# Oxygen transport parameter in plasma membrane of eye lens fiber cells by saturation recovery EPR

**DOI:** 10.1101/2020.05.28.121285

**Authors:** N. Stein, W. K. Subczynski

## Abstract

A probability distribution of rate constants contained within an exponential-like saturation recovery (SR) electron paramagnetic resonance signal can be constructed using stretched exponential function fitting parameters. Previously (Stein et al. *Appl. Magn. Reson.* 2019.), application of this method was limited to the case where only one relaxation process, namely spin-lattice relaxations due to the rotational diffusion of the spin labels in the intact eye-lens membranes, contributed to an exponential-like SR signal. These conditions were achieved for thoroughly deoxygenated samples. Here, the case is described where the second relaxation process, namely Heisenberg exchange between the spin label and molecular oxygen that occurs during bimolecular collisions, contributes to the decay of SR signals. We have further developed the theory for application of stretched exponential function to analyze SR signals involving these two processes. This new approach allows separation of stretched exponential parameters, namely characteristic stretched rates and heterogeneity parameters for both processes. Knowing these parameters allowed us to separately construct the probability distributions of spin-lattice relaxation rates determined by the rotational diffusion of spin labels and the distribution of relaxations induced strictly by collisions with molecular oxygen. The later distribution is determined by the distribution of oxygen diffusion concentration products within the membrane, which forms a sensitive new way to describe membrane fluidity and heterogeneity. This method was validated *in silico* and by fitting SR signals from spin-labeled intact nuclear fiber cell plasma membranes extracted from porcine eye lenses equilibrated with different fractions of air.

**Statement of Significance:** Multi-exponential spin-lattice relaxation in complex membranous systems can be described by a stretched exponential function that provides a continuous probability distribution of relaxation rates rather than discreet relaxations from separate domains. The stretched exponential function has two fitting parameters, the characteristic spin-lattice relaxation rate (T_1str_^−1^) and the stretching parameter (β), obtained without any assumption about the number of membrane domains and their homogeneity. For membranes equilibrated with air, collisions with molecular oxygen provide an additional relaxation pathway for spin labels that depends on the oxygen-diffusion-concentration product in the vicinity of spin labels. This new approach allows separation of membrane fluidity and heterogeneity sensed by motion of lipid spin labels from those described by the translational diffusion of molecular oxygen.

## Introduction

Molecular oxygen is an effective probe molecule for studying membrane lipid packing. Molecular oxygen is small, hydrophobic, and fast diffusing; therefore, it can enter even transiently formed, small vacant pockets in a lipid bilayer membrane. Because of that, the oxygen diffusion-concentration product reflects the dynamics of *gauche-trans* isomerization of acyl chains and structural nonconformability of neighboring lipids (1–3). Because lipid packing is affected by the lipid composition and the presence of membrane proteins, the solubility and diffusion of molecular oxygen are different in induced membrane domains. Due to the paramagnetic nature of molecular oxygen, a dual-probe saturation recovery (SR) electron paramagnetic resonance (EPR) method allows observation of the bimolecular collision rate between spin labels and molecular oxygen that depends on the local oxygen-diffusion-concentration product, termed oxygen transport parameter (OTP) (4).

First developed in the 1980s, this oximetry method was used to calculate the permeability coefficient for oxygen across model (1, 5–7) and biological (8–11) membranes. Comparing this coefficient with the permeability coefficient of a water layer of the same thickness as the membrane allowed determination of whether the membranes form a barrier to oxygen transport. The logical conclusion of such a finding was that membrane domains of different compositions must have different OTP profiles and, thus, different oxygen permeability coefficients. Spin-label oximetry has since been used to differentiate bulk, boundary, and trapped lipids (9, 12–14); liquid ordered phase domains (15–17); and pure cholesterol bilayer domains (16–22) within model and biological membranes.

The SR EPR spin labeling method measures the spin-lattice relaxation rate (*T*_1_^−1^) of nitroxide spin labels in membranes, which, in magnetically diluted samples, is determined by the rotational diffusion of the probe (23–27). In the dual-probe SR EPR method, molecular oxygen (a fast-relaxing paramagnetic species) induces spin exchange during collisions with spin labels (a slow-relaxing paramagnetic species), which provides an additional relaxation pathway for spin labels. It provides a quantitative and the most sensitive way to measure the collision rate between a spin label and molecular oxygen (8, 28).

The SR signal is recorded as an exponential-like decay and can be analyzed using mono- and multi-exponential decay functions. Due to the nonorthogonal nature of exponential function, the fitting requires previous knowledge or a strong assumption about the number of exponentials and their values. In membrane studies, it means that the number of membrane domains should be preliminary defined. Due to the practical limitation of fitting, this number is limited to two or three homogeneous domains. This is convenient for describing model membranes; however, it is problematic for describing complex biological membrane samples. These samples may consist of mixtures of membranes with different lipid and protein compositions. For example, the composition of an eye lens membrane changes gradually as cells mature (29, 30, 39–42, 31–38). These changes induce variability in domains and local environments.

The application of the stretched exponential function (SEF) to analyze SR EPR signals from spin-labeled biological membranes removes the requirement for a specific number of discrete exponentials. It reduces the number of fitting parameters to two: the characteristic spin-lattice relaxation rate (*T*_1str_^−1^) and the heterogeneity parameter *β*. When *β* is 1, the function is a single exponential and the environment is homogeneous. When *β* is less than one, more than one exponential is present. The fitting parameters provide the means to construct a probability distribution of rates contained within the signal.

Previously, we used the SEF to analyze SR data obtained from deoxygenated intact cortical and nuclear fiber cell plasma membranes extracted from porcine eye lenses and spin-labeled with phospholipid and cholesterol analogs (43). The constructed probability distributions of spin-lattice relaxation rates due to the rotational diffusion provides quantitative information about the rigidity and heterogeneity of the samples. The method of separating the OTPs and spin-lattice relaxation rates due to the rotational diffusion when fitting the data to distinct exponentials was also reported previously (4). Here, we develop a method to separate spin-lattice relaxation rates due to the rotational diffusion and oxygen collision in the complex biological samples where assumption of the numbers and values of distinct exponentials is not desirable or feasible. The term *stretched oxygen transport parameters (SOTP)* is defined, and two methods for its extraction from the total SR EPR signal are outlined. These methods are evaluated *in silico* and with real data from complex biological membranes.

## Outline of theory

### Stretched exponential as a sum of exponential decays

The SEF can be used to describe a distribution of decay rate constants (*k*) as a sum of exponential decays (44):

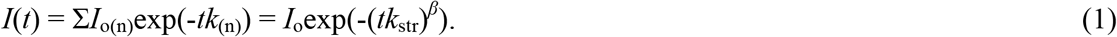

In this equation, *I*(t) is the signal amplitude at time (*t*), *I*_o(n)_ is a fractional contribution of each exponential term n toward the signal at time zero, *I*_o_ is a normalized signal amplitude at time zero, *k*_(n)_ is the individual decay rate constant, *k*_str_ is a characteristic or stretched rate constant, and *β* is the heterogeneity parameter that corresponds to *k*_str_.

The distribution of rate constants can be generated using the two fitting parameters obtained from the fitting of the multi-exponential-like signal to the SEF, the characteristic rate constant, *k*_str_, and the *β*, as shown in Fig. 1. This distribution can be used to evaluate the probability of finding a range of rate constants within the signal and, in that, the portion of the signal that may arise from the range of these constants.

**Figure 1.**
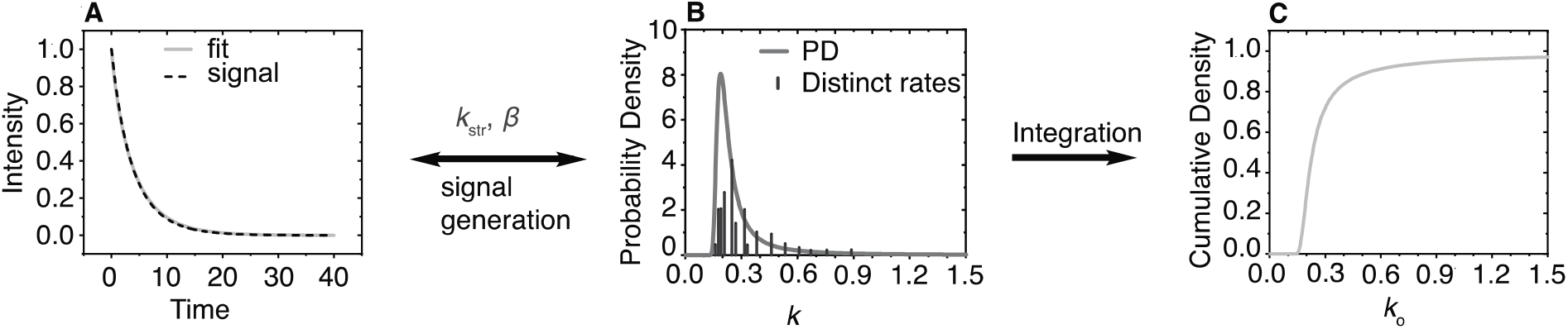
Illustration of the principle of using the SEF as a sum of exponentials. In *(A)*, the dashed line is a signal generated from the sum of 15 random exponential decay constants, which are represented as the vertical lines in *(B)*. The relative heights of these lines reflect their fractional contribution to the signal, and the position the value of the rate constant. The solid line in (A) represents the SEF of the signal with two parameters, *k*_str_ and *β*. These parameters were used to construct the continuous probability distribution as described in Stein *et al.* (43). This distribution covers all the rate constants within the signal. *(C)* The cumulative distribution function was obtained by integrating the area under the probability distribution and can be used to assess the probability of finding a range of rate constants within the signal.

The *β* parameter determines the shape of the distribution; hence, it is termed *heterogeneity parameter*. When *k*_str_ is multiplied by a factor, all individual rates within the distribution range are multiplied by that same factor; see Fig. 2. Conversely, if all rates in a distribution range are multiplied by the same factor, as is the case in membrane oximetry, the *k*_str_ will increase by that same factor and the beta parameter will remain the same. This relationship of the SEF parameters to probability distribution is key to extracting the distribution of OTPs using data obtained by SR EPR.

**Figure 2.**
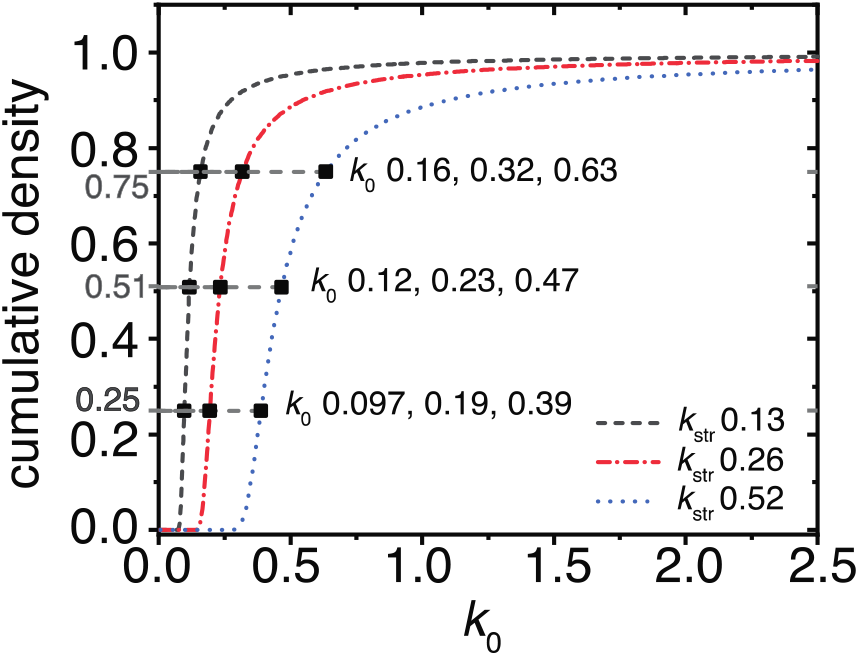
Illustration of stretched rate constant properties. The cumulative density of three stretched exponential distributions that possess the same *β* parameter (0.90), and *k*_str_ parameters that differ by the factor of two, as indicated. Consequently, the *k*_o_s at cumulative density values of 0.25, 0.51, and 0.75 increase by the factor of two. Hence, as rates within the distribution are multiplied by a factor of two, so is the *k*_str_.

### Oxygen transport parameter

In SR EPR, the OTP, which is defined as the oxygen diffusion-concentration product, is measured as the effective oxygen collision rate with spin labels (4). In the magnetically diluted samples (under nitrogen atmosphere), the SR signal (mainly) depends on the rotational diffusion of the spin probe. If no heterogeneity exists within the rotational diffusion of the probes, the signal can be fitted with a single exponential decay:

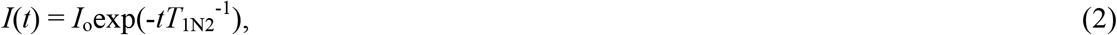

where *T*_1N2_^−1^ is the rotational diffusion spin-lattice relaxation rate.

Because molecular oxygen is paramagnetic, the collision between molecular oxygen and a spin label provides an additional relaxation pathway for the spin label. The increase in the relaxation rate of the signal can be described by the product of two exponentials:

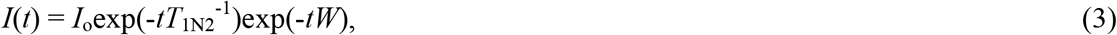

where *W* is the effective oxygen collision rate with spin labels in samples saturated with air at 760 mmHg.

If no heterogeneity in *W* exists, then the signal can be fitted to a single exponential:

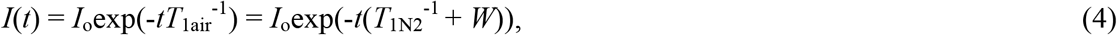

where *T*_1Air_^−1^ is the spin-lattice relaxation in samples saturated with air.

*W* is the difference between *T*_1Air_^−1^ and *T*_1N2_^−1^ (4):

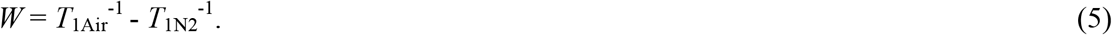

In other words, *W* is the slope of the straight line formed by the observed spin-lattice relaxation rates (*T*_1obs_^−1^s) at various air fractions:

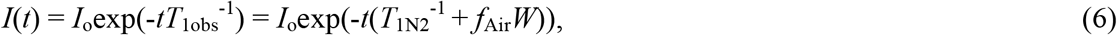

where *f*_Air_ is the air fraction at which the sample is equilibrated before the signal is recorded.

### Stretched oxygen transport parameters

#### Special case

When the SR EPR signal is recorded in the presence of molecular oxygen, two independent processes contribute to the spin-lattice relaxation process: rotational diffusion and oxygen collision. Therefore, in heterogenous samples, these rates can be described by two distributions of relaxation rates.

If the spin-lattice relaxation is homogenous under nitrogen atmosphere but the oxygen transport parameter is not, the signal will consist of the sum of products of exponentials in Eq. 7a. The *T*_1N2_^−1^ can be readily factored out as in Eq. 7b.

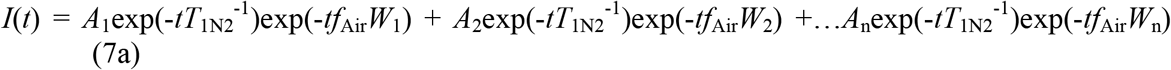

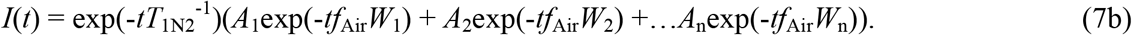

The multi-exponential portion can be replaced with the stretched exponential term, providing the SOTP:

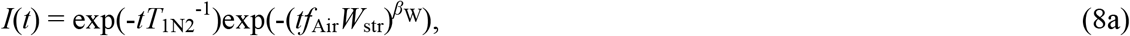

where *W*_str_ is the stretched oxygen collision rate and *β*_W_ is the heterogeneity of the *W*_str_.

Eq. 8a represents the special case where the SOTP is fitted directly while holding the rotational diffusion parameter obtained under the nitrogen constant. *W*_str_ is extracted either from the slope of *f*_Air_*W*_str_ versus air fraction or by fitting the *W*_str_ of a sample equilibrated in air. *β*_W_ represents the heterogeneity of the SOTP term and does not change with air fraction. In biological membranes, rotational diffusion term *T*_1N2_^−1^ is rarely homogenous. The relaxation processes by the rotational diffusion or Heisenberg exchange are independent as are the distributions of the rates by which they are represented. Because of that, the special case also can be applied to instances where the rotational diffusion measured under nitrogen is heterogenous:

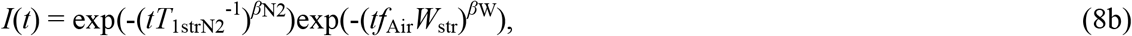

where *β*_N2_ represents the heterogeneity of the rotational diffusion spin-lattice relaxation rates, and *T*_1strN2_^−1^ is the characteristic rotational diffusion spin-lattice relaxation rates. Because the addition of oxygen does not change these parameters, the SOTP can be fitted directly when holding parameters obtained under the nitrogen constant.

The special case relies on the data obtained in the absence of oxygen for fitting.

#### General case

We developed an alternative, “general case” to extract SOTP parameters from the total signal, which does not use the rotational diffusion spin-lattice relaxation parameter.

At any air fraction, the observed relaxation rate (*T*_1strobs_^−1^) represents the sum of the products of the rotational diffusion spin-lattice relaxation rate, and the OTP is multiplied by the air fraction at which the observation was recorded:

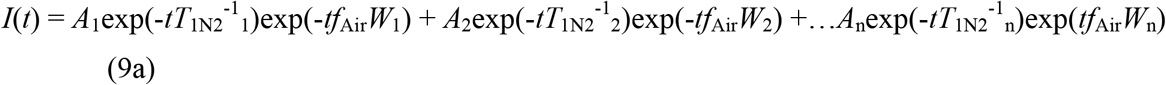

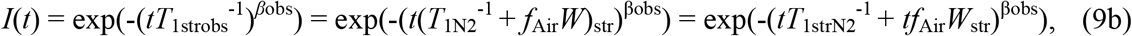

where the heterogeneity of that term is *β*_obs_. During air titration, the rate only increases when the probe collides with molecular oxygen; therefore, W_str_ can be extracted from the slope of the straight line formed by *T*_1strobs_^−1^s versus air fraction. The y-intercept of such line corresponds to *T*_1strN2_^−1^.

More difficult is determining the observed heterogeneity behavior. In the absence of air (*i.e.*, when *f*_Air_*W*_str_ is zero), the observed heterogeneity corresponds to that of the rotational diffusion spin-lattice relaxation rates. When the *f*_Air_*W*_str_ term is sufficiently large (*i.e.*, when *tT*_1strN2_^−1^ is negligible by comparison), the *β*_obs_ corresponds to the heterogeneity of the OTPs. The behavior of *β*_obs_ between the two extremes was determined empirically and can be described as the weighted sum of the heterogeneities (*β* parameters) contributed by the rotational diffusion and oxygen collision rates. Eq. 10, which describes this dependence, was validated *in silico* (see Fig. 3) and through the fitting the experimental data to Eq. 10.

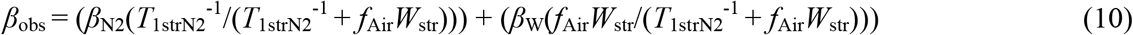

**Figure 3.**
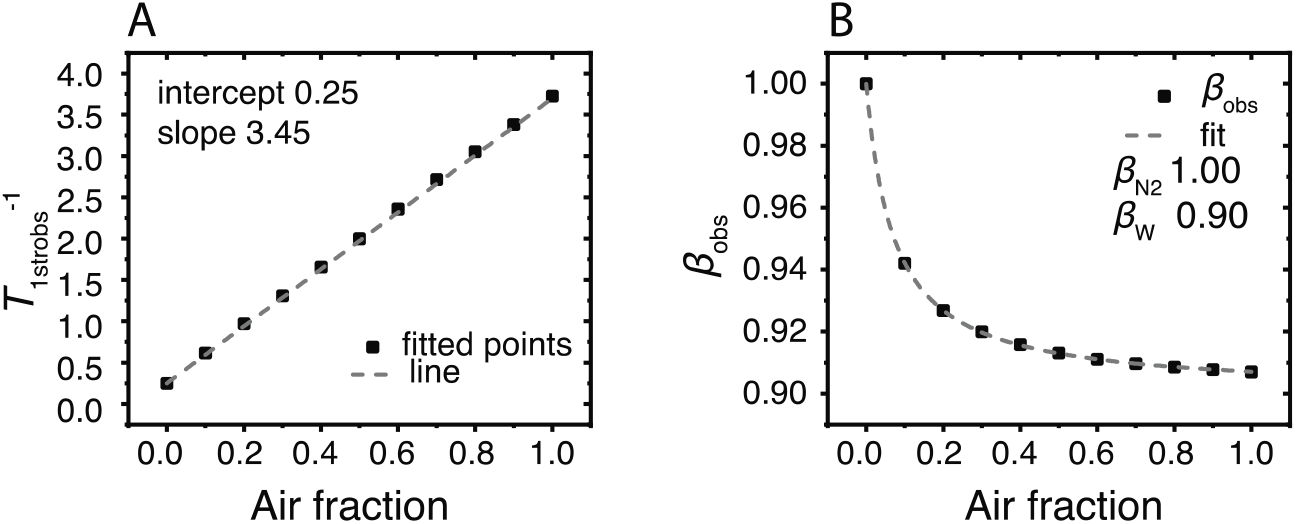
Signals that were simulated using Eq. 8a with the parameters *T*_1N2_^−1^ 0.25, *W*_str_ 3.45, *β*_W_ 0.90, and various *f*_Air_ were fitted with Eq. 9b. The fitted parameters were then plotted versus the air fraction. *(A)* shows *T*_1strobs_^−1^ versus air fraction, and *(B)* shows *β*_obs_ versus air fraction. In (A), the square dots are the fitted values and the dashed line is the linear fit of the data. In (B), the square dots are the fitted values and the dashed line is the fit with Eq. 10; the *W*_str_ and T_1N2_^−1^ parameters obtained from the linear fit in (A) were held constant.

## Materials and Methods

### Materials

The spin-labeled cholesterol analog (androstane spin label [ASL]) was purchased from Molecular Probes (Eugene, OR); see the structures in Fig. 1 of Ref. (18). Other chemicals of at least reagent grade were purchased from Sigma-Aldrich (St. Louis, MO).

### Isolation of nuclear fiber cells

The following procedures were described previously (14, 45). Porcine eyes from 2-year-old animals were obtained on the day of slaughter from **Johnsonville Sausage**, LLC (Watertown, WI). Lenses were dissected and kept at −80°C until needed. Next, the lenses were thawed and decapsulated, and the nuclear portion of the fiber cells was separated from the cortical portion based on their consistencies (37, 46) and used in experiments.

### Isolation of intact fiber cell membranes

Intact nuclear membranes from 36 eye lenses were isolated from tissue, as reported previously (47–49), using the method developed in (50) with minor modifications. Special care was taken to produce a uniform suspension by repeatedly aspirating the solution through a syringe fitted with an 18-gauge needle. The final pellet was washed and resuspended in 10 mM PIPES pH 7.0, 150 mM NaCl. Half of the suspension was stored at −80°C; the other half was washed with water and lyophilized.

### Isolation of total lipids

The total lipids were extracted from the lyophilized membranes using the Folch procedure (51) with minor modifications. The resultant lipid samples were soft, white solids and were stored at −20°C.

### Spin labeling of intact membranes

Spin labeling was performed as described previously (45). ASL film was prepared on the bottom of a test tube by drying the appropriate amount of spin label in chloroform (to achieve the molar ratio of spin label/total lipids of ~1/100 in each sample). Intact membrane suspensions (~0.2 mL containing 1 to 2 mg of total lipids) were added to the test tubes and shaken for about 2 h at room temperature. This incubation time was enough to incorporate all the spin-label molecules into the membranes. Finally, membrane suspensions were centrifuged for a short time, and the loose pellet was transferred to a 0.6 mm i.d. capillary made of gas-permeable methylpentene polymer (*i.e.*, TPX) and used for EPR measurements (52).

### Preparation of lens lipid membranes

Multilamellar vesicles made from the total lipids contained 1 mol% of ASL. They are referred to as “lens lipid membranes (LLMs)” and were prepared using the rapid solvent exchange method (53–55). The final membrane dispersions (1 to 2 mg of total lipids) were centrifuged briefly (12,000 g, 15 min, 4°C), and the loose pellet was used for EPR measurements.

### SR EPR measurements

The SR EPR signal for each sample was obtained on the central line of the ASL EPR spectrum by short-pulse SR EPR at X-band using a loop-gap resonator (1, 56). The signal was recorded 100 ns (dead time) after a short (300 ns duration) saturating pulse of 1 W power. All measurements of samples equilibrated with the same gas that was used for temperature control (*i.e.*, a controlled mixture of nitrogen and dry air adjusted with flowmeters [model 7631H-604; Matheson Gas Products, Irvine, TX]) were carried out at 36°C (52). The record time was at least 20 times the stretched relaxation time of the sample, and the sampling interval was approximately 1/80^th^ of the stretched relaxation time.

### Analysis of SR signals using the SEF

All data fitting was performed using the OriginPro Version 2019 OriginLab Corporation (Northampton, MA) software package. The SR signals were fitted as appropriate with Eqs. 8a, 8b, and 9b using the Levenberg-Marquardt algorithm. The zero time was adjusted to 220 ns after the end of the pulse. When Eq. 8a or 8b was used, the *T*_1_^−1^ and *β*_N2_ values obtained from fitting the anaerobic sample with the single exponential were held constant as *T*_1N2_^−1^. When Eq. 10 was used to fit *β*_obs_ versus air fraction, the *T*_1strN2_^−1^ and *W*_str_ values were held constant while the *β*_N2_ and *β*_W_ values were free to vary.

The probability densities were computed in Matlab R2018a (MathWorks, Natick, MA) using the previously reported probability distribution function (44) and adjusted for either *T*_1strN2_^−1^ or *W*_str_ as described in (43). Additionally, a cutoff time for high rates was implemented at 22.7 μs^−1^. Due to our time delay, rates faster than that would have been at less than 1% of their initial amplitude.

## Results and Discussion

### Special and general cases: A comparison of samples with homogenous T_1N2_^−1^s in LLMs

Here, we compare SOTP (*W*_str_s and *β*_W_s) and rotational diffusion (*T*_1strN2_^−1^ and *β*_N2_) parameters extracted using Eqs. 8a (special case) and 9b (general case) from experimentally obtained SR signals of ASL labeled LLMs. The results are summarized in Fig. 4.

**Figure 4.**
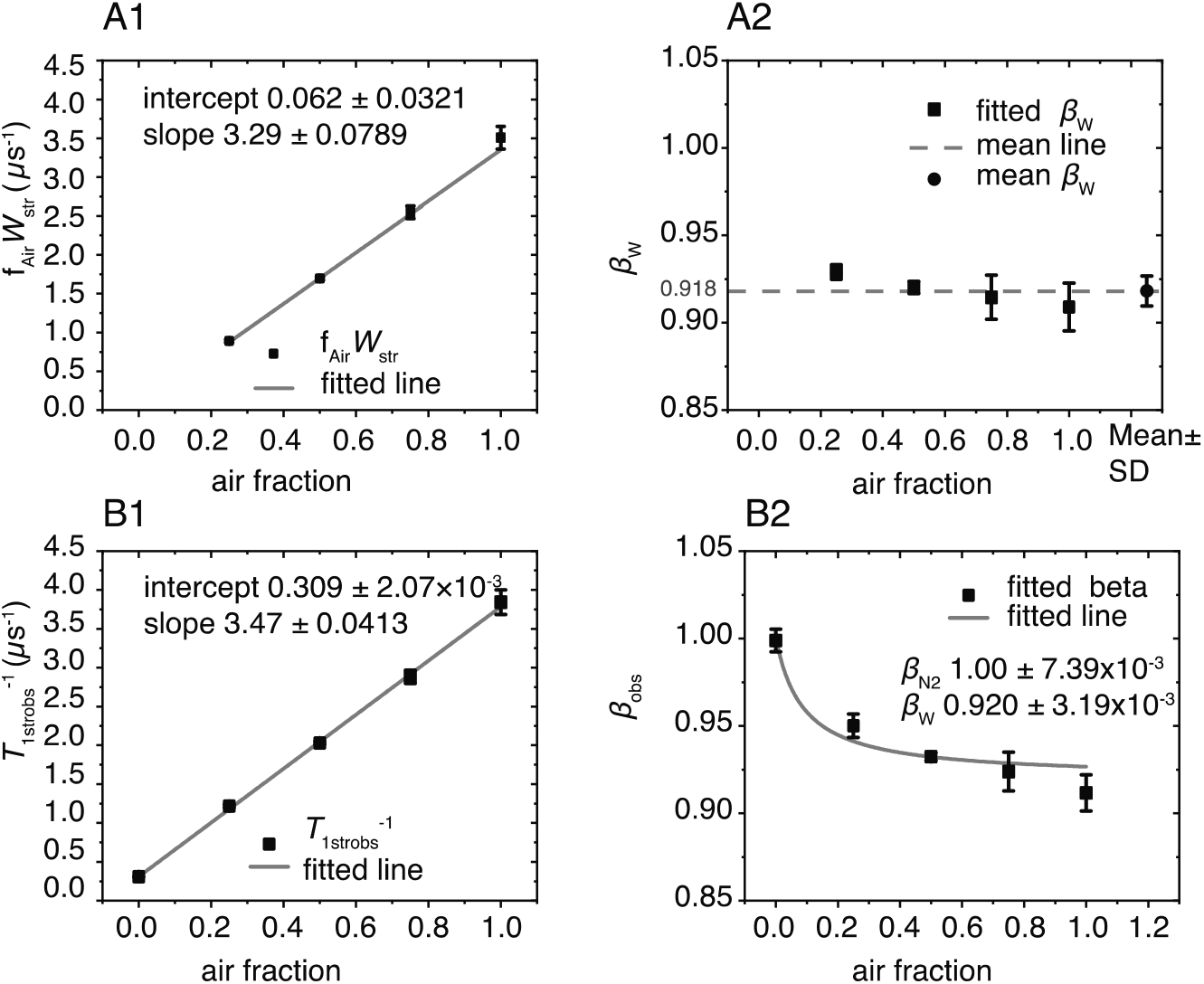
The fitting parameters *f*_Air_*W*_str_, *β*_W_, *T*_1strobs_^−1^, and *β*_obs_ obtained from fitting the SR signals of the LLM using appropriate equations and plotted versus the air fractions at which they were obtained. *(A1)* The average *f*_Air_*W*_str_ values obtained from fitting at least three SR signals using Eq. 8a are indicated by black squares with vertical bars that represent the standard deviations. The gray line is the linear fit of these data points with an intercept of 0.062 ± 0.032 and a slope of 3.29 ± 0.0789 μs^−1^. The values following the ± sign are standard errors of the parameters. *(A2)* The black squares are the average values of *β*_Ws_ obtained from fitting at least three SR signals using Eq. 8a. The vertical bars represent their standard deviations. The average of these values was found to be 0.918 ± 8.55 × 10^−3^ and is represented by a black circle; the standard deviations are represented by vertical bars, respectively. The dashed gray line indicates where the measured values fall in relation to the mean. *(B1)* The average *T*_1strobs_^−1^ values obtained from fitting at least three SR signals using Eq. 9b are indicated by black squares with vertical bars that represent the standard deviations. The gray line is the linear fit of these data points with an intercept of 0.309 ± 2.07 × 10^−3^ and a slope of 3.29 ± 0.0413 μs^−1^. The values following the ± sign are standard errors of the parameters. *(B2)* The black squares are the average values of β_obs_ obtained from at least three SR signals, and the vertical lines correspond to standard deviations. The fit to Eq. 10 is indicated by the gray line. The values of the intercept and slope obtained from the linear fit of the points in (B1) represent *T*_1strN2_^−1^ and *W*_str_ respectively, and were held constant during fitting. The β_N2_ and β_W_ parameters were allowed to vary and converged at 1.00 ± 7.38 × 10^−3^ and 0.920 ± 3.19 × 10^−3^, respectively. The values following the ± signs are the standard errors of the parameters.

When fitted with Eq. 9b, the averaged *T*_1strobs_^−1^ and *β*_obs_ values from at least three SR signals obtained anaerobically were 0.309 ± 1.65 × 10^−3^ μs^−1^ and 1.00 ± 6.37 × 10^−3^, respectively. The value of *β*_obs_ indicates that the spin-lattice relaxation rate is a single exponential and that rotational diffusion of the spin labels is homogenous. However, as demonstrated in Fig. 4 (A2 and B2), the LLMs contain more than one OTP.

*W*_str_ was obtained from the slope of a straight line formed by either *f*_Air_*W*_str_ or *T*_1strobs_^−1^ versus air fraction Fig. 4 (A1and B1). *W*_str_ extracted from the slope of the straight line formed by the values obtained from Eq. 8a (*f*_Air_*W*_str_ versus air fraction) was 3.29 ± 7.89 × 10^−2^ μs^−1^, and that extracted from the slope of the line formed by the values obtained from Eq. 9b (*T*_1strobs_^−1^ versus air fraction) was 3.47 ± 4.12 × 10^−2^ μs ^−1^. These values have an approximately 5% difference.

To check for self-consistency, the y-intercept of each graph also was analyzed. When Eq. 8a was used the obtained y-intercept, −6.22 × 10^−2^ ± 3.21 × 10^−2^ μs^−1^ was close to its theoretical value of zero. When Eq. 9b was used, the y-intercept of the *T*_1strobs_^−1^ versus air fraction line matched the value obtained under nitrogen 0.309 ± 4.08 × 10^−2^ μs^−1^.

The *β*_W_ values obtained from Eq. 8a are plotted versus the air fraction that they were obtained at in Fig. 4A2. Their average was found to be 0.918 ± 8.55 × 10^−3^. When using Eq. 9b, the *β*_W_ of 0.920 ± 3.19 × 10^−3^ and *β*_N2_ of 1.00 ± 7.39 × 10^−3^ were elucidated by fitting the graph of *β*_obs_ versus air fraction with Eq. 10. The corresponding *T*_1strN2_^−1^ and *W*_str_ values obtained from the rate graph Fig. 4B1 were held constant. Notably, the fitted *β*_N2_ matches the *β*_obs_ obtained anaerobically of 1.00 ± 6.37 × 10^−3^ and confirms the self-consistency of the results.

Together, these analyses demonstrate that Eqs. 8a and 9b provide comparable and self-consistent rate and heterogeneity parameter values. The rate parameter variability of 5% may be due to the noise. The heterogeneity parameters display a similar pattern of being higher than the fitted value at lower air fractions and lower than the fitted value at higher air fractions. These deviations may be due to the spectrometer noise and higher number of averages required to obtain a signal at higher air fraction.

### Special and general cases: A comparison of samples with heterogenous T_1N2_^−1^s in intact membranes

To demonstrate the application of “special” and “general” cases to analysis of the SR signals from samples that showed heterogeneity under deoxygenated conditions, we isolated intact porcine nuclear eye lens membranes and labeled them with ASL. The SR signals for samples equilibrated at different air fractions were obtained, as described in the Methods section. These signals were fitted using Eqs. 8b and 9b, and the results are summarized in Fig. 5.

**Figure 5.**
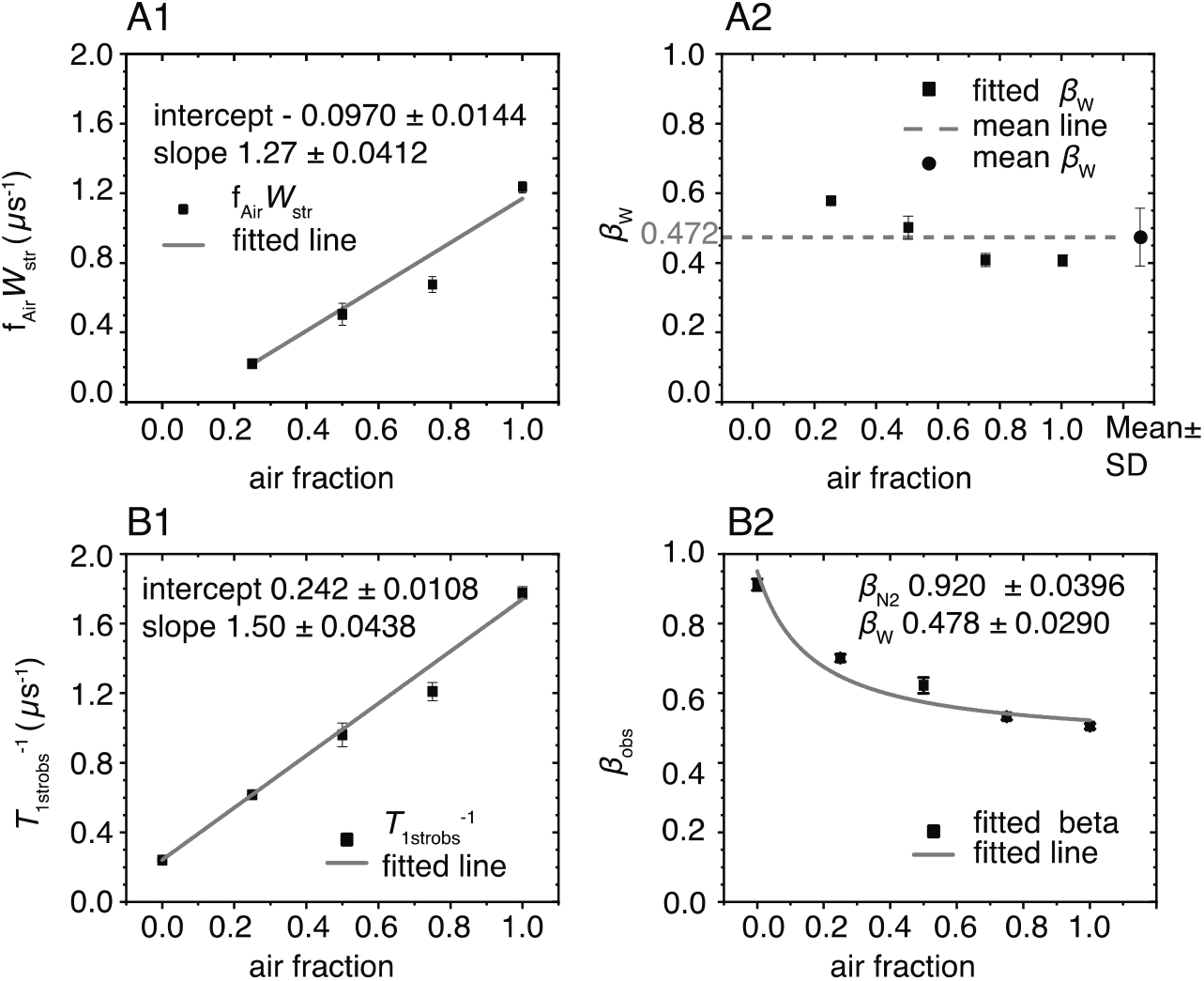
The fitting parameters *f*_Air_*W*_str_, *β*_W_, *T*_1strobs_^−1^, and *β*_obs_ obtained from air titration of intact nuclear eye lens membranes plotted versus the air fractions at which the signals were obtained. *(A1)* The black squares are the average *f*_Air_*W*_str_ values obtained from fitting at least three SR signals using Eq. 8b. The vertical bars represent the corresponding standard deviations. The gray line is the linear fit of these data points with an intercept of −9.70×10^−2^ ± 1.44×10^−2^ and a slope of 1.27 ± 4.12 ×10^−2^ μs^−1^. The values following the ± sign are standard errors of the parameters. *(A2)* The black squares are the average values of *β*_Ws_ obtained from fitting at least three SR signals using Eq. 8b. The vertical bars represent their standard deviations. The black circle represents the average of these values of 0.472, and the vertical bars indicate the standard deviation of 0.0766. The dashed gray line indicates where the measured values fall in relation to the mean. *(B1)* The average *T*_1strobs_^−1^ values obtained from fitting at least three SR recovery 1strobs signals using Eq. 9b are indicated by black squares with vertical bars that represent the standard deviations. The gray line is the linear fit of these data points with an intercept 0.242 ± 1.08 × 10^−2^ and a slope of 1.50 ± 4.38 × 10^−2^ μs^−1^. The values following the ± sign are standard errors of the parameters. *(B2)* The black squares are the average values of *β*_obs_ obtained from at least three SR signals, and the vertical lines correspond to standard deviations. The fit to Eq. 10 is presented by the gray line. The values of the intercept and slope obtained from the linear fit of the points in (B1) represent *T*_1strN2_^−1^ and *W*_str_, respectively, and were held constant during fitting. The *β*_N2_ and β_W_ parameters were allowed to vary and converged at 0.920 ± 0.0396 and 0.477 ± 0.0290, respectively. The values following the ± signs are the standard errors of the parameters.

The average *T*_1strN2_^−1^ and the corresponding standard deviation of at least three SR signals obtained under nitrogen was found to be 0.241 ± 6.94 × 10^−3^ μs^−1^ with a corresponding *β*_N2_ of 0.912 ± 0.0167. To extract SOTP parameters, these values were held constant while the signals were fit to Eq. 8b. The fitted *f*_Air_*W*_str_ values were plotted against the air fractions at which they were obtained in Fig. 5A1. The slope of the straight line formed by these points, which corresponds to *W*_str_, was found to be 1.27 ± 4.12 × 10^−2^ μs^−1^. Similarly, the *T*_1strobs_^−1^ values obtained from fitting the same signals to Eq. 9b were plotted versus the air fractions at which they were obtained, and *W*_str_ of 1.50 ± 4.38 × 10^−2^ μs^−1^ was obtained from the slope of the straight line formed by these points. Although the errors of fitting are comparable, the resulting two values differ from one another by 15%.

We evaluated the y-intercepts of each graph to check the self-consistency of results. The “special case” rate line, which should pass through the origin, came in at −9.70 × 10^−2^ ± 1.44×10^−2^. The “general case” line was 0.242 ± 1.08×10^−2^ μs^−1^ and matches the *T*_1strN2_^−1^ obtained under nitrogen.

The heterogeneity parameter *β*_W_ for the special case was obtained directly by fitting the signals using Eq 8b. The average of these values was 0.472 ± 7.66 × 10^−2^. This value is close 0.478 ± 2.90 × 10^−2^, which is the value obtained by fitting the *β*_obs_ versus air fraction graph with Eq. 10, as demonstrated in Fig. 5B2. The *W*_str_ and *T*_1strN2_^−1^ were kept constant during the fitting. According to that fit, the *β*_N2_ is 0.920 ± 3.99 × 10^−2^ and is close to the value of 0.912 ± 1.67 × 10^−2^ obtained under nitrogen (Fig. 5).

The SR signals from the intact membrane are noisier than those from liposomes. The measurements may contain a larger error that is reflected in the slight inconsistency of results obtained by the “special case” and “general case.”

### Distribution of spin-lattice relaxation rates of ASL in intact membranes and LLMs due to rotational diffusion

To make ultimate sense of the fitting parameters obtained by the SEF, we constructed and analyzed probability distributions (43, 44). First, as demonstrated in Fig. 6, we did so for the parameters obtained for membranes equilibrated with nitrogen, when they reflected the rotational diffusion of lipid spin labels. Spin-lattice relaxation rates due to the rotational diffusion are heterogenous in intact membranes, while in LLMs the SR signal is fitted by a single exponential function with the spin-lattice relaxation rate depicted by a single line (delta function). The distribution of rates in intact membranes indicates that 81% of the signal is due to relaxation rates slower than those of the LLMs, and the remaining are faster.

**Figure 6.**
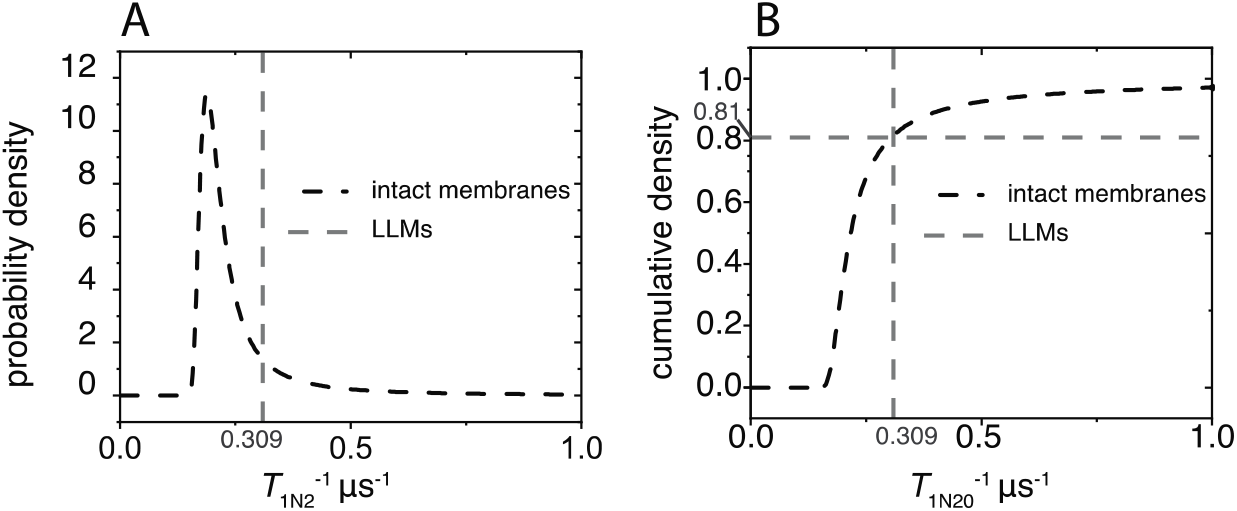
The probability distribution densities of rotational diffusion spin-lattice relaxation rates and the integrals (cumulative densities) were constructed as described in the text. *(A)* The probability distribution density for the spin-lattice relaxation rates associated with the rotational diffusion of ASL in the intact membrane is represented by the dashed black line. The dashed gray line at 0.309 μs^−1^ represents the rotational diffusion spin-lattice relaxation rate of ASLs in LLMs. *(B)* The dashed black line is the cumulative density of rotational diffusion spin-lattice relaxation rates for ASL in the intact membranes. The gray dashed line signifies the rotational diffusion spin-lattice relaxation rate of ASLs in LLMs. The crossing point between these lines indicates that 81% of the signal in the intact membranes is from ASLs that relaxed slower than those in the LLMs, and the remainder of the signal relaxed faster.

LLMs approximate bulk lipid domains in intact membranes and are expected to be the most fluid in terms of rotational diffusion. The presence of faster spin-lattice relaxation rates due to rotational diffusion in the intact membranes can be explained by the fact that the LLMs are composed of lipids from all layers in the nuclear region that are pulled together and made into multilamellar vesicles. The lipids in the intact membranes maintain their unique lipid compositions for each isolated fiber cell layer, and the SR signal is the sum of signals from membranes of each layer. Additionally, intact membranes possess integral and peripheral proteins that induce specific lipid organization and asymmetry (57–59) that may be responsible in a more fluid environments than in LLMs.

### Distribution of OTPs in intact membranes and LLMs

The extracted SOTP parameters were used to construct distribution of OTPs in intact membranes and in LLMs as outlined in (43). The distributions in Fig. 7, allowed us to make the following observations. The LLMs exhibit OTP heterogeneity that indicates the presence of various domains. The heterogeneity of the OTPs in intact membranes is significantly larger than that in LLMs. In intact membranes, 66% of the collisions of oxygen with spin labels possess collision rates slower than 2.23 μs^−1^. Fewer than 0.01% account for such slow rates in LLMs. The slow rates may be indicative of trapped lipid domains where oxygen concentration is low, or diffusion is slow. The crossing point of cumulative density curves in Fig. 7B indicates that, similar to rotational diffusion, a higher density of fast oxygen collision rates is present in the intact membranes than in LLMs. This correlation indicates that the organization of intact membranes in the sample is considerably more heterogenous than in the LLM sample.

**Figure 7.**
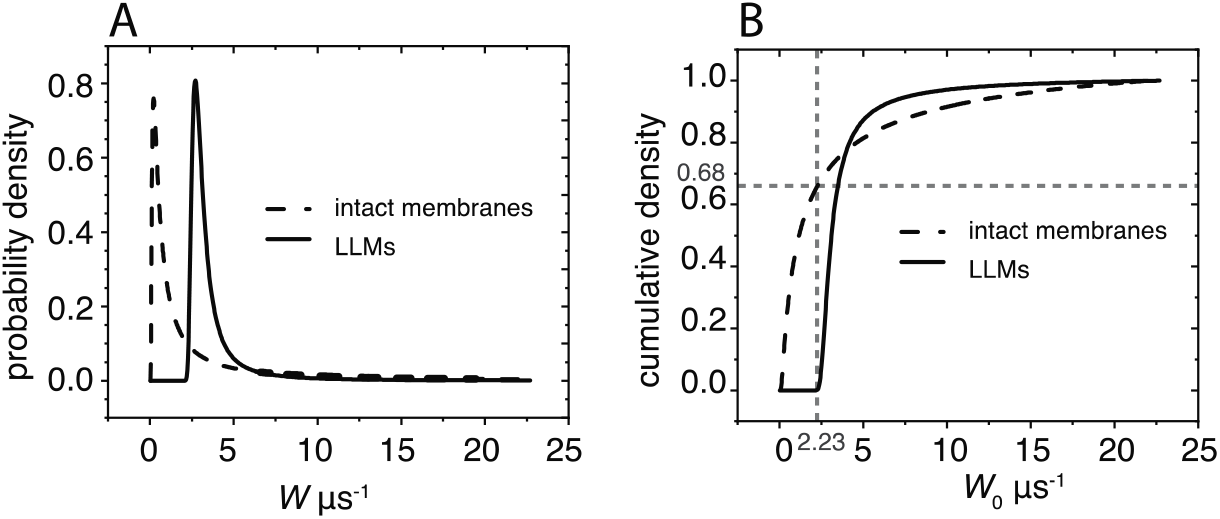
The probability density distributions of OTPs sensed by ASL in LLMs and intact membranes and their integrals were constructed as described in the text. *(A)* The probability density distributions of OTPs in intact membranes are indicated by a dashed black line, and in LLMs by a solid black line. *(B)* The integrals of these distributions are the cumulative densities. The dashed black lines are the cumulative densities of OTPs in intact membranes, and the solid black line is that of LLMs. The gray dashed lines indicate that 68% of the density in the intact membranes falls below 2.23 μs^−1^, whereas less than 0.01% of the density of the LLMs can be found below that rate.

## Conclusion

SR EPR data from spin labeled lipid membranes provide membrane physical properties in terms of the rotational diffusion of lipids and oxygen-diffusion-concentration product within the membrane. In our previous publication (43), using SEF parameters, we constructed the probability distributions of spin-lattice relaxation rates due to the rotational diffusion in the biological spin-labeled membrane samples coming from porcine eye lenses. This data treatment removes the need to fit a complex sample to the distinct exponentials. Because cholesterol, phospholipid (29, 33, 42, 46), and protein compositions (48, 60) of fiber cell membranes change gradually as the cells age, and our samples are composed of many layers of such cells, the number of different environments experienced by spin labels may be difficult to estimate. The SEF method of analysis allows us to consider the minute physical property differences in membranes of cells from different layers as well as the different domains within those membranes. Our understanding of the properties of the SEF fitting parameters allowed us to take on a new level of complexity when two independent relaxation processes contribute to SR signals. The additional relaxation pathway was provided by paramagnetic molecules of oxygen that, through collisions with spin labels, contribute to the decay of the SR signals. To the best of our knowledge, we are the first to apply the SEF analysis to a system in which two relaxation processes contribute to the exponential-like decay signal. We developed theory to analyze SR signals with the SEF, which allowed us to obtain stretched exponential fitting parameters separately for each contributing process and applied it to practice.

Because any enhancement of the spin-lattice relaxation rate during air titration is due to Heisenberg exchange with oxygen, the “special” case method (Eqs. 8a and 8b) relies on accurately obtained rotational diffusion parameters. These remain constant at any oxygen partial pressure so long as other experimental conditions remain constant. Holding rotational diffusion parameters constant allows to extract SOTP parameters *W*_str_ and *β*_W_ at any known oxygen concentration.

The “general” case relies on observation that the product of two stretched exponential terms produces a relaxation curve that can be fitted to a third stretched exponential. The stretched rate of the new exponential was found to correspond to the sum of the stretched rates of the two contributing processes (Eq. 9b). Thus, analogous to the previously defined OTP (4), the rate component of SOTP is defined as:

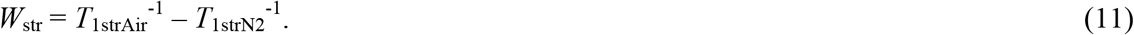

A *β* parameter remains constant when all rates within a distribution are multiplied by the same factor. The *β*_obs_ of a product of the two stretched exponentials is the sum of the two parameters weighted by their stretched exponential rates (Eq. 10).

We demonstrated that the advanced theory for the application of the SEF to analyze SR signals for membranes saturated with molecular oxygen allowed construction of (a) probability distributions of spin-lattice relaxation rates determined by the rotational diffusion of spin labels, and (b) the distribution of relaxations induced strictly by collisions with molecular oxygen. These probability distributions were constructed for simple LLMs and intact nuclear fiber cell membranes. They show the effects of integral and peripheral membrane proteins on membrane fluidity, as well as the heterogeneity of membrane fluidity sensed by the cholesterol analog spin label alone (Fig. 6) and sensed by the movement and solubility of molecular oxygen (Fig. 7). The latter distribution is determined by the distribution of oxygen-diffusion-concentration products within the membrane, which forms a flexible new way to describe membrane fluidity and heterogeneity. Use of lipid spin labels, SR EPR, stretched exponential analysis, and discrimination by oxygen transport offer a powerful approach to analyze complex membranous samples derived from biological tissue.

## Author Contributions

Conceptualization, NS, WKS; performed research, NS; data analysis, NS; writing - original draft preparation, NS; review & editing, WKS; visualization, NS; project administration, WKS; funding acquisition, WKS.

## Acknowledgements

Research reported in this publication was supported by the National Eye Institute of the National Institutes of Health under award number R01 EY015526. The content is solely the responsibility of the authors and does not necessarily represent the official views of the National Institutes of Health.

## References

1. Subczynski, W.K., J.S. Hyde, A. Kusumi, W.K. Subczynski, and A. Kusumi. 1991. Effect of Alkyl Chain Unsaturation and Cholesterol Intercalation on Oxygen Transport in Membranes: A Pulse ESR Spin Labeling Study. Biochemistry. 30:8578–8590.

2. Pace, R.J., and S.I. Chan. 1982. Molecular motions in lipid bilayers. I. Statistical mechanical model of acyl chain motion. J. Chem. Phys. 76:4217–4227.

3. Altenbach, C., D.A. Greenhalgh, H.G. Khorana, and W.L. Hubbell. 1994. A collision gradient method to determine the immersion depth of nitroxides in lipid bilayers: Application to spin-labeled mutants of bacteriorhodopsin. Proc. Natl. Acad. Sci. U. S. A. 91:1667–1671.

4. Kusumi, A., W.K. Subczynski, and J.S. Hyde. 1982. Oxygen transport parameter in membranes as deduced by saturation recovery measurements of spin-lattice relaxation times of spin labels. Proc. Natl. Acad. Sci. U. S. A. 79:1854–1858.

5. Subczynski, W.K., J.S. Hyde, and A. Kusumi. 1989. Oxygen permeability of phosphatidylcholine-cholesterol membranes. Proc. Natl. Acad. Sci. U. S. A. 86:4474–4478.

6. Widomska, J., M. Raguz, and W.K. Subczynski. 2007. Oxygen permeability of the lipid bilayer membrane made of calf lens lipids. Biochim. Biophys. Acta - Biomembr. 1768:2635–2645.

7. Subczynski, W.K., and J.S. Hyde. 1998. Membranes: Barriers or Pathways for Oxygen Transport. In: Hudetz AG, editor. Oxygen Transport to Tissue XX, Advances in Experimental Medicine and Biology. Plenum, New York: . pp. 399–408.

8. Subczynski, W.K., L.E. Hopwood, and J.S. Hyde. 1992. Is the mammalian cell plasma membrane a barrier to oxygen transport? J. Gen. Physiol. 100:69–87.

9. Raguz, M., L. Mainali, W.J. O’Brien, and W.K. Subczynski. 2015. Lipid domains in intact fiber-cell plasma membranes isolated from cortical and nuclear regions of human eye lenses of donors from different age groups. Exp. Eye Res. 132:78–90.

10. Subczynski, W.K., and H.M. Swartz. 2005. EPR Oximetry in Biological and Model Samples. in Biological Magnetic resonance. In: Eaton SS, GR Eaton, editors. Vol. 23. Biomedical ESR - Part A: Free Radicals, Metals, Medicine, and Physiology. Kluwer, Boston: . pp. 229–282.

11. Subczynski, W.K., J. Widomska, and L. Mainali. 2017. Factors determining the oxygen permeability of biological membranes: Oxygen transport across eye lens fiber-cell plasma membranes: Oxygen transport across eye lens fiber-cell plasma membranes. In: Halpern H, JC LaManna, DK Harrison, B Epel, editors. Oxygen Transport to Tissue XXXIX. Advances in Experimental Medicine and Biology Vol. 977. Springer. pp. 27–34.

12. Ashikawa, I., J.J. Yin, W.K. Subczynski, J.S. Hyde, T. Kouyama, and A. Kusumi. 1994. Molecular Organization and Dynamics in Bacteriorhodopsin-Rich Reconstituted Membranes: Discrimination of Lipid Environments by the Oxygen Transport Parameter Using a Pulse ESR Spin-Labeling Technique. Biochemistry. 33:4947–4952.

13. Kawasaki, K., J.J. Yin, W.K. Subczynski, J.S. Hyde, and A. Kusumi. 2001. Pulse EPR detection of lipid exchange between protein-rich raft and bulk domains in the membrane: Methodology development and its application to studies of influenza viral membrane. Biophys. J. 80:738–48.

14. Raguz, M., L. Mainali, W.J. O’Brien, and W.K. Subczynski. 2014. Lipid-protein interactions in plasma membranes of fiber cells isolated from the human eye lens. Exp. Eye Res. 120:138–151.

15. Subczynski, W.K., A. Wisniewska, J.S. Hyde, and A. Kusumi. 2007. Three-dimensional dynamic structure of the liquid-ordered domain in lipid membranes as examined by pulse-EPR oxygen probing. Biophys. J. 92:1573–1584.

16. Raguz, M., L. Mainali, J. Widomska, and W.K. Subczynski. 2011. The immiscible cholesterol bilayer domain exists as an integral part of phospholipid bilayer membranes. Biochim. Biophys. Acta - Biomembr. 1808:1072–1080.

17. Subczynski, W.K., M. Raguz, and J. Widomska. 2010. Studying lipid organization in biological membranes using liposomes and EPR spin labeling. Methods Mol. Biol. 606:247–269.

18. Raguz, M., L. Mainali, J. Widomska, and W.K. Subczynski. 2011. Using spin-label electron paramagnetic resonance (EPR) to discriminate and characterize the cholesterol bilayer domain. Chem. Phys. Lipids. 164:819–829.

19. Mainali, L., M. Raguz, and W.K. Subczynski. 2013. Formation of cholesterol bilayer domains precedes formation of cholesterol crystals in cholesterol/dimyristoylphosphatidylcholine membranes: EPR and DSC studies. J. Phys. Chem. B. 117:8994–9003.

20. Subczynski, W.K., M. Raguz, J. Widomska, L. Mainali, and A. Konovalov. 2012. Functions of cholesterol and the cholesterol bilayer domain specific to the fiber-cell plasma membrane of the eye lens. J. Membr. Biol. 245:51–68.

21. Mainali, L., M. Pasenkiewicz-Gierula, and W.K. Subczynski. 2020. Formation of cholesterol Bilayer Domains Precedes Formation of Cholesterol Crystals in Membranes Made of the Major Phospholipids of Human Eye Lens Fiber Cell Plasma Membranes. Curr. Eye Res. 45:162–172.

22. Widomska, J., W.K. Subczynski, L. Mainali, and M. Raguz. 2017. Cholesterol Bilayer Domains in the Eye Lens Health: A Review. Cell Biochem. Biophys. 75:387–398.

23. Mailer, C., R.D. Nielsen, and B.H. Robinson. 2005. Explanation of spin-lattice relaxation rates of spin labels obtained with multifrequency saturation recovery EPR. J. Phys. Chem. A. 109:4049–4061.

24. Marsh, D. 2018. Molecular order and T1-relaxation, cross-relaxation in nitroxide spin labels. J. Magn. Reson. 290:38–45.

25. Mainali, L., J.B. Feix, J.S. Hyde, and W.K. Subczynski. 2011. Membrane fluidity profiles as deduced by saturation-recovery EPR measurements of spin-lattice relaxation times of spin labels. J. Magn. Reson. 212:418–425.

26. Mainali, L., J.S. Hyde, and W.K. Subczynski. 2013. Using spin-label W-band EPR to study membrane fluidity profiles in samples of small volume. J. Magn. Reson. 226:35–44.

27. Robinson, B.H., D.A. Haas, and C. Mailer. 1994. Molecular dynamics in liquids: Spin-lattice relaxation of nitroxide spin labels. Science (80-.). 263:490–493.

28. Subczynski, W.K., and J.S. Hyde. 1981. The diffusion-concentration product of oxygen in lipid bilayers using the spin-label T1 method. BBA - Biomembr. 643:283–291.

29. Huang, L., V. Grami, Y. Marrero, D. Tang, M.C. Yappert, V. Rasi, and D. Borchman. 2005. Human lens phospholipid changes with age and cataract. Investig. Ophthalmol. Vis. Sci. 46:1682–1689.

30. Paterson, C.A., J. Zeng, Z. Husseini, D. Borchman, N.A. Delamere, D. Garland, and J. Jimenez-Asensio. 1997. Calcium ATPase activity and membrane structure in clear and cataractous human lenses. Curr. Eye Res. 16:333–338.

31. Truscott, R.J.W. 2000. Age-related nuclear cataract: A lens transport problem. Ophthalmic Res. 32:185–194.

32. Yappert, M.C., M. Rujoi, D. Borchman, I. Vorobyov, and R. Estrada. 2003. Glycero-versus sphingo-phospholipids: Correlations with human and non-human mammalian lens growth. Exp. Eye Res. 76:725–734.

33. Borchman, D., W.C. Byrdwell, and M.C. Yappert. 1994. Regional and age-dependent differences in the phospholipid composition of human lens membranes. Investig. Ophthalmol. Vis. Sci. 35:3938–3942.

34. Deeley, J.M., T.W. Mitchell, X. Wei, J. Korth, J.R. Nealon, S.J. Blanksby, and R.J.W. Truscott. 2008. Human lens lipids differ markedly from those of commonly used experimental animals. Biochim. Biophys. Acta - Mol. Cell Biol. Lipids. 1781:288–298.

35. Rujoi, M., R. Estrada, and M.C. Yappert. 2004. In Situ MALDI-TOF MS Regional Analysis of Neutral Phospholipids in Lens Tissue. Anal. Chem. 76:1657–1663.

36. Raguz, M., J. Widomska, J. Dillon, E.R. Gaillard, and W.K. Subczynski. 2009. Physical properties of the lipid bilayer membrane made of cortical and nuclear bovine lens lipids: EPR spin-labeling studies. Biochim. Biophys. Acta - Biomembr. 1788:2380–2388.

37. Rujoi, M., J. Jin, D. Borchman, D. Tang, and M.C. Yappert. 2003. Isolation and lipid characterization of cholesterol-enriched fractions in cortical and nuclear human lens fibers. Investig. Ophthalmol. Vis. Sci. 44:1634–1642.

38. Zelenka, P.S. 1984. Lens lipids. Curr. Eye Res. 3:1337–1359.

39. Bassnett, S., Y. Shi, and G.F.J.M. Vrensen. 2011. Biological glass: Structural determinants of eye lens transparency. Philos. Trans. R. Soc. B Biol. Sci. 366:1250–1264.

40. Gonen, T., Y. Cheng, J. Kistler, and T. Walz. 2004. Aquaporin-0 membrane junctions form upon proteolytic cleavage. J. Mol. Biol. 342:1337–1345.

41. Kistler, J., and S. Bullivant. 1980. Lens gap junctions and orthogonal arrays are unrelated. FEBS Lett. 111:73–78.

42. Vidová, V., J. Pól, M. Volný, P. Novák, V. Havlícek, S.K. Wiedmer, and J.M. Holopainen. 2010. Visualizing spatial lipid distribution in porcine lens by MALDI imaging high-resolution mass spectrometry. J. Lipid Res. 51:2295–2302.

43. Stein, N., L. Mainali, J.S. Hyde, and W.K. Subczynski. 2019. Characterization of the Distribution of Spin-Lattice Relaxation Rates of Lipid Spin Labels in Fiber Cell Plasma Membranes of Eye Lenses with a Stretched Exponential Function. Appl. Magn. Reson. 50:903–918.

44. Johnston, D.C. 2006. Stretched exponential relaxation arising from a continuous sum of exponential decays. Phys. Rev. B - Condens. Matter Mater. Phys. 74:184430.

45. Mainali, L., M. Raguz, W.J. O’Brien, and W.K. Subczynski. 2013. Properties of fiber cell plasma membranes isolated from the cortex and nucleus of the porcine eye lens. Exp. Eye Res. 182:1432–1440.

46. Estrada, R., and M.C. Yappert. 2004. Regional phospholipid analysis of porcine lens membranes by matrix-assisted laser desorption/ionization time-of-flight mass spectrometry. In: Journal of Mass Spectrometry. . pp. 1531–1540.

47. Cenedella, R.J., and C.R. Fleschner. 1992. Selective association of crystallins with Lens “native” membrane during dynamic cataractogenesis. Curr. Eye Res. 11:801–815.

48. Chandrasekher, G., and R.J. Cenedella. 1995. Protein associated with human lens “native” membrane during aging and cataract formation. Exp. Eye Res. 60:707–717.

49. Lim, J., Y.C. Lam, J. Kistler, and P.J. Donaldson. 2005. Molecular characterization of the cystine/glutamate exchanger and the excitatory amino acid transporters in the rat lens. Investig. Ophthalmol. Vis. Sci. 46:2869–2877.

50. Bloemendal, H., A. Zweers, F. Vermorken, I. Dunia, and E.L. Benedetti. 1972. The plasma membranes of eye lens fibres. Biochemical and structural characterization. Cell Differ. 1:91–106.

51. Folch, J., M. Lees, and G.H. Sloane Stanley. 1957. A simple method for the isolation and purification of total lipides from animal tissues. J. Biol. Chem. 226:497–509.

52. Subczynski, W.K., C.C. Felix, C.S. Klug, and J.S. Hyde. 2005. Concentration by centrifugation for gas exchange EPR oximetry measurements with loop-gap resonators. J. Magn. Reson. 176:244–248.

53. Buboltz, J.T. 2009. A more efficient device for preparing model-membrane liposomes by the rapid solvent exchange method. Rev. Sci. Instrum. 80:124301.

54. Huang, J., J.T. Buboltz, and G.W. Feigenson. 1999. Maximum solubility of cholesterol in phosphatidylcholine and phosphatidylethanolamine bilayers. Biochim. Biophys. Acta - Biomembr. 1417:89–100.

55. Mainali, L., M. Raguz, W.J. O’Brien, and W.K. Subczynski. 2013. Properties of membranes derived from the total lipids extracted from the human lens cortex and nucleus. Biochim. Biophys. Acta - Biomembr. 1828:1432–1440.

56. Mainali, L., T.G. Camenisch, J.S. Hyde, and W.K. Subczynski. 2017. Saturation Recovery EPR Spin-Labeling Method for Quantification of Lipids in Biological Membrane Domains. Appl. Magn. Reson. 48:1355–1373.

57. Arumugam, S., E.P. Petrov, and P. Schwille. 2015. Cytoskeletal pinning controls phase separation in multicomponent lipid membranes. Biophys. J. 108:1104–1113.

58. Corradi, V., B.I. Sejdiu, H. Mesa-Galloso, H. Abdizadeh, S.Y. Noskov, S.J. Marrink, and D.P. Tieleman. 2019. Emerging Diversity in Lipid-Protein Interactions. Chem. Rev. 119:5775–5848.

59. Fujimoto, T., and I. Parmryd. 2017. Interleaflet coupling, pinning, and leaflet asymmetry-major players in plasma membrane nanodomain formation. Front. Cell Dev. Biol. 4:155.

60. Thibault, D.B., C.J. Gillam, A.C. Grey, J. Han, and K.L. Schey. 2008. MALDI Tissue Profiling of Integral Membrane Proteins from Ocular Tissues. J. Am. Soc. Mass Spectrom. 19:814–822.

